# Regulation of transcription factor SP1 by β-catenin destruction complex modulates Wnt response

**DOI:** 10.1101/308841

**Authors:** Rafeeq Mir, Ankita Sharma, Saurabh J. Pradhan, Sanjeev Galande

## Abstract

The ubiquitous transcription factor Specificity protein 1 (SP1) is heavily modified post-translationally. These modifications are critical for switching its functions and modulation of its transcriptional activity, DNA-binding and stability. However the mechanism governing the stability of SP1 by cellular signaling pathways is not well understood. Here, we provide biochemical and functional evidences that SP1 is an integral part of the Wnt signaling pathway. We identified a phosphodegron motif in SP1 that is specific to mammals. In absence of Wnt signaling, GSK3β kinase mediated phosphorylation and β-TrCP E3 ubiquitin ligase mediated ubiquitination is required to induce SP1 degradation. When Wnt signaling is on, SP1 is stabilized in β-catenin-dependent manner. SP1 directly interacts with β-catenin and Wnt signaling induces the stabilization of SP1 by impeding its interaction with β-TrCP and AXIN1, components of the destruction complex. Wnt signaling suppresses ubiquitination and subsequent proteosomal degradation of SP1. Furthermore, SP1 regulates Wnt-dependent stability of β-catenin and their mutual stabilization is critical for target gene expression, suggesting a feedback mechanism. Upon stabilization SP1 and β-catenin co-occupy the promoters of TCFL2/β-catenin target genes. Collectively, this study uncovers a direct link between SP1 and β-catenin in Wnt signaling pathway.

## Introduction

Wnt signaling plays essential role in tissue homeostasis, cell fate determination, proliferation and embryonic development (1, 2). Mutations in Wnt signaling pathway are associated with developmental defects and cancers (1, 3–5). The core event in the Wnt pathway is stabilization of β-catenin upon activation of Wnt signaling. The stabilization of β-catenin is regulated by cytoplasmic destruction complex consisting of scaffold protein AXIN1, APC, GSK3β and CSK1α. In absence of Wnt ligand (WNT OFF condition) β-catenin is engaged by destruction complex followed by sequential phosphorylation by CSK1α and GSK3β (6, 7). Phosphorylated β-catenin is recognized by the ubiquitin E3 ligase β-TrCP and subsequently degraded by proteosomal pathway. In presence of Wnt ligand (WNT ON condition) β-catenin is disengaged from cytoplasmic destruction complex by multivesicular sequestering of GSK3β and other mechanisms. Thereby β-catenin evades phosphorylation and recognition by β-TrCP and hence degradation by proteosomal pathway. Dephosphorylated β-catenin translocates inside nucleus, interacts with the LEF/TCF transcription factors to drive target gene expression (2, 7). Wnt/β-catenin target genes play critical role in tumor development and progression. The components of Wnt signaling cascade play critical role in tumor development and poor prognosis, hence can be targeted for therapeutic intervention in cancers driven by dysregulation of Wnt pathway (3).

Specificity protein1 (SP1), a member of SP family proteins containing C2H2 zinc finger domain (8) was earlier thought to be ubiquitously expressed and involved in transcriptional regulation of housekeeping genes. However, multiple recent evidences suggest tissue-specific role of SP1 in regulating genes considered as hallmarks of cancers and genes required during development and differentiation (9, 10). SP1 is heavily decorated by various post-translational modifications such as O-linked glycosylation, acetylation, phosphorylation, ubiquitination and sumoylation that are critical for switching SP1 functions. Number of signaling kinases have been implicated in SP1 post-translational modification that influence SP1’s mode of action, DNA-binding, stability and transactivation (10). Considering the role of SP1 in regulating gene expression required during embryonic development and progression of cancers (9–13), it is essential to understand the mechanism regulating SP1 expression. Although phosphorylation mediated degradation of SP1 at protein level is reported (14,15), how phosphorylation takes place in cellular context and thereby affects its stability is not fully elucidated.

Aberrant activation of Wnt signaling has been implicated in various cancers mediated by mutations in various regulators of this pathway (16). Recently, crosstalk between novel targets and hyperactivation of Wnt signaling has been implicated in regulating the fate of Wnt signaling (17, 18) Our earlier study further provided evidences that this crosstalk with novel targets plays critical role in regulating colorectal tumorigenesis and regulation of Wnt responsive genes (19). Interestingly, both SP1 and β-catenin possess the regulatory phosphodegron motif and interact with β-TrCP and thereby hinting at possible overlap of regulation of β-catenin and SP1. Overlap of regulation and function between β-catenin and SP1 has been reported (14, 20). Thus, based on these studies, there is a possibility of SP1 and β-catenin molecular crosstalk regulated by a common pathway and hence regulating the Wnt target genes.

Here we report that SP1 is a downstream target of Wnt signaling pathway. We show that β-catenin is required for SP1 stability in GSK3β-dependent manner. We found that the cytoplasmic destruction complex required for β-catenin destabilization is also required for SP1 destabilization. We show that Wnt signaling induces stabilization of SP1 by impeding its interaction with the E3 ubiquitin ligase β-TrCP and AXIN1. We present a direct link demonstrating that SP1 regulates the stability of β-catenin by impeding its interaction with cytosolic destruction complex. Further, destabilization of the destruction complex and mutual stabilization of SP1 and β-catenin is required for Wnt/β-catenin signaling dependent regulation of Wnt responsive genes.

## Results

### Wnt signaling promotes SP1 stabilization

To elucidate the regulation of SP1, we analyzed the expression of SP1 in Wnt signaling driven colorectal cancer cells in comparison with primary cell line CRL1790. The expression pattern indicates that Wnt driven cancer cells exhibit elevated levels of SP1 (Figure 1A). To test whether stimulation of Wnt signaling could be involved in regulation of SP1, we treated HEK293 cells with Wnt3A. The activation of Wnt signaling induced the expression of SP1 (Figure 1B). Similarly, L3 cells showed significant upregulation of SP1 along with β-catenin upon activation of Wnt pathway using Wnt conditioned medium (Figure 1D and 1E). Since HEK293 cells have intact destruction complex and respond to Wnt stimulation, we sought to use these cells to determine whether Wnt stimulation by Wnt ligand Wnt3A can induce stability of SP1 at protein level. We thus treated HEK293 FLAG-SP1 expressing cells with Wnt3A in time-dependent manner (scheme for hypothesis presented in Figure 1C). The treatment of FLAG-SP1 HEK 293 cells induced the robust stability of SP1 at protein level as determined by levels of FLAG tag in comparison with control (Figure 1F, compare lane 2 with lane 3 and lane 4). The Wnt stimulation also induced the stability of known target β-catenin indicating the activation of Wnt signaling cascade. Similarly, Wnt3A treatment in FLAG-SP1 expressing L3 cells induced stabilization of SP1 (Figure 1G, compare lanes 2 and 3, Supplementary Figure 1A). This analysis revealed that Wnt pathway activation is required to induce the stability of SP1 and thus could be the reason that Wnt driven colorectal cancer cells express higher levels of SP1. Next, to determine if Wnt stimulation induces the stability of SP1 in cytosol and nucleus, we analyzed expression of FLAG tagged SP1 in cytosolic and nuclear fractions. Immunoblot analysis revealed that Wnt stimulation induces SP1 both in cytosolic and nuclear fractions (Figure 1H, compare lanes 2 and 3, 4 and 5). Next, to investigate that stability of SP1 requires inhibition of proteosomal pathway, we treated FLAG-SP1 expressing HEK293 cells with proteosomal pathway inhibitor MG132. Immunoblot analysis using anti- FLAG antibody indicated that inhibition of proteosomal pathway induced the stability of SP1 (Figure 1I).

**Figure 1:**
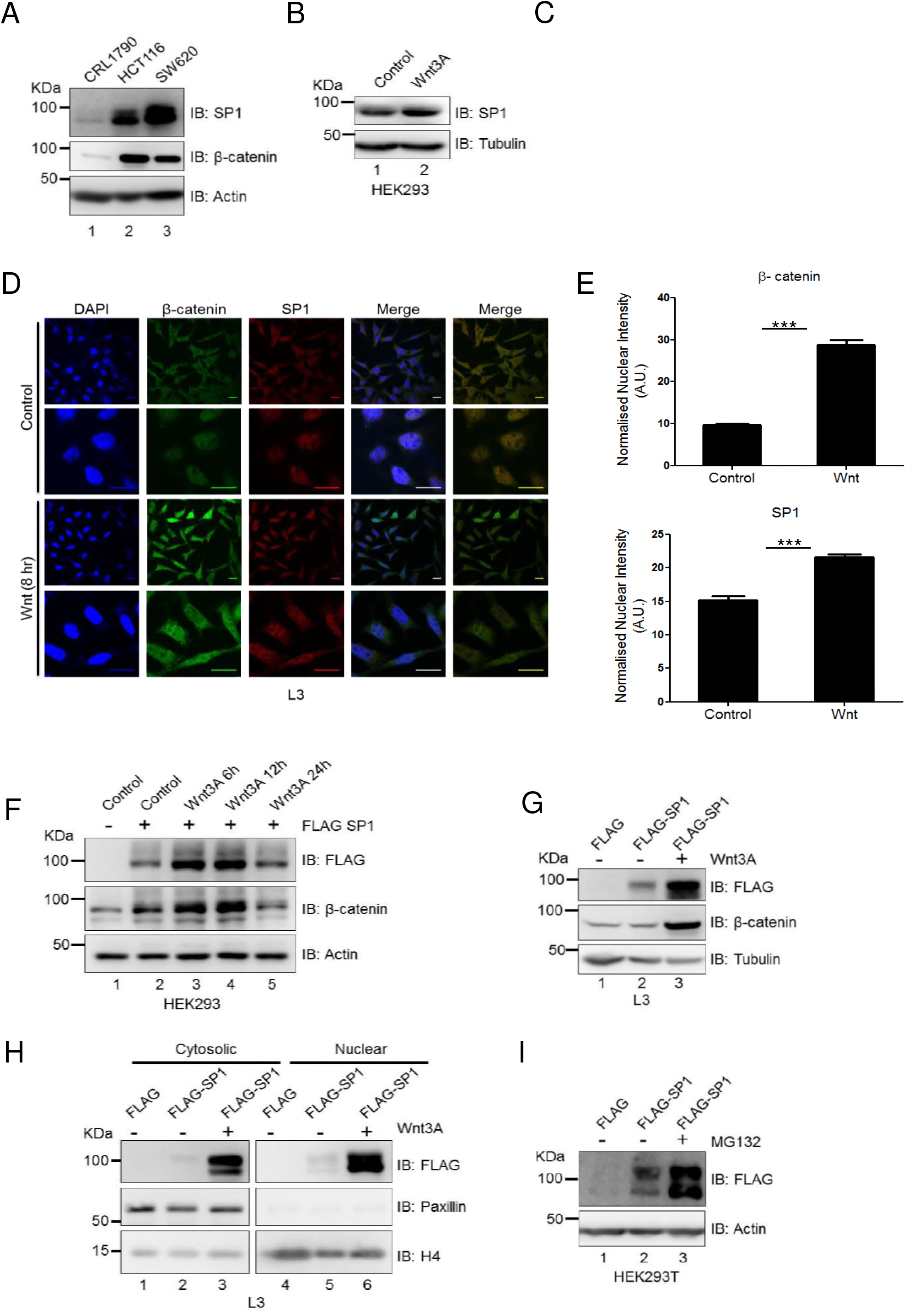
Wnt/β-catenin signaling induces SP1 stabilization. **(A)** Immunoblot for expression of SP1 and β-catenin in CRL1790, HCT116 and SW620. **(B)** Immunoblot for expression of SP1 in L3 mouse epithelial cells treated with Wnt3A for 6h. **(C)** Schematic depicting probable mode of Wnt signaling mediated stabilization of SP1. **(D)** Immunofluorescence assay showing increase in cytosolic and nuclear levels of SP1 and β-catenin upon Wnt stimulation in L3 cells. (Scale bar represents 20 μm). First merged panel indicates merging of all three chanels whereas in the second merged panel the DAPI is removed to better visualize co-localization of nuclear β-catenin and SP1. **(E)** Quantification of nuclear intensities of β-catenin and SP1 levels (Mann-Whitney test, Two tailed, P<0.0001). **(F)** Immunoblot for FLAG and β-catenin in FLAG SP1 expressing HEK293 cells after treatment with Wnt3A in time-dependent manner. **(G)** Immunoblot for FLAG and β-catenin in FLAG-SP1 expressing L3 cells upon treatment with Wnt3A. Tubulin was used as endogenous control. **(H)** Immunoblot for FLAG, Paxillin and Histone H4 in cytosolic and nuclear fractions of FLAG-SP1 expressing L3 cells upon treatment with Wnt3A. **(I)** Immunoblot for FLAG-SP1 in control and MG132 treated cells. HEK293 cell were transfected with FLAG-SP1 and treated with DMSO and proteosome pathway inhibitor MG132 for 4 h.

### SP1 interacts with β-catenin in colorectal cancer cells

Next, to determine whether SP1 physically interacts with β-catenin, we performed co-immunoprecipitation assay in HCT-15 colorectal cancer cells. Immunoblot analysis following co-immunoprecipitation revealed that SP1 interacts with β-catenin (Figure 2A, lane 3 and lane 4). Such interaction was also observed in COLO205 wherein immunoprecipitation with anti-β-catenin pulled down SP1 (Figure 2B). Next, we tested the validity of data in different cellular model; we overexpressed mutant S37A stabilized form of β-catenin in HeLa cells and immunoprecipitated with anti- β-catenin antibody. The analysis reveals that SP1 physically interacts with β-catenin (Figure 2C). To further confirm if SP1 interacts with β-catenin, we performed immunofluorescence assay in L3 Wnt3A cells, which have constitutively active Wnt signalling. The data revealed that SP1 co-localizes with β-catenin (Figure 2D). To determine whether SP1 interacts with β-catenin directly, we performed *in vitro* GST pull-down assays using FLAG-SP1 expressing HEK293 cells (Figure 2E, lane 3). Similar results were observed in GST pull-down assay with GST-SP1 using lysates of FLAG β-catenin expressing HEK293 cells (Figure 2F, lane 3) and HCT116 cells for interaction with endogenous SP1 (Supplementary Figure S2B left panel, lane 3). These observations confirmed that SP1 and β-catenin interact directly in vitro. Further, to delineate which domain of β-catenin interacts with SP1 we performed pull-down assays using various GST-tagged domains of β-catenin (Figure 2G). GST pull-downs revealed that both N- and C-termini of β-catenin interact with SP1 but not the arm domain (Figure 2H, lane 3 and lane 5). Further, both wild type SP1 and delta phosphodegron mutant SP1 interact with β-catenin with equal efficiency (Supplementary Figure S2C and S2D). Furthermore, GST-SP1 pull-down assay revealed that SP1 interacts with β-catenin through its N-terminus (Figure 2I). Collectively, these findings suggest that SP1, stabilized upon Wnt/β-catenin signaling, interacts with β-catenin.

**Figure 2:**
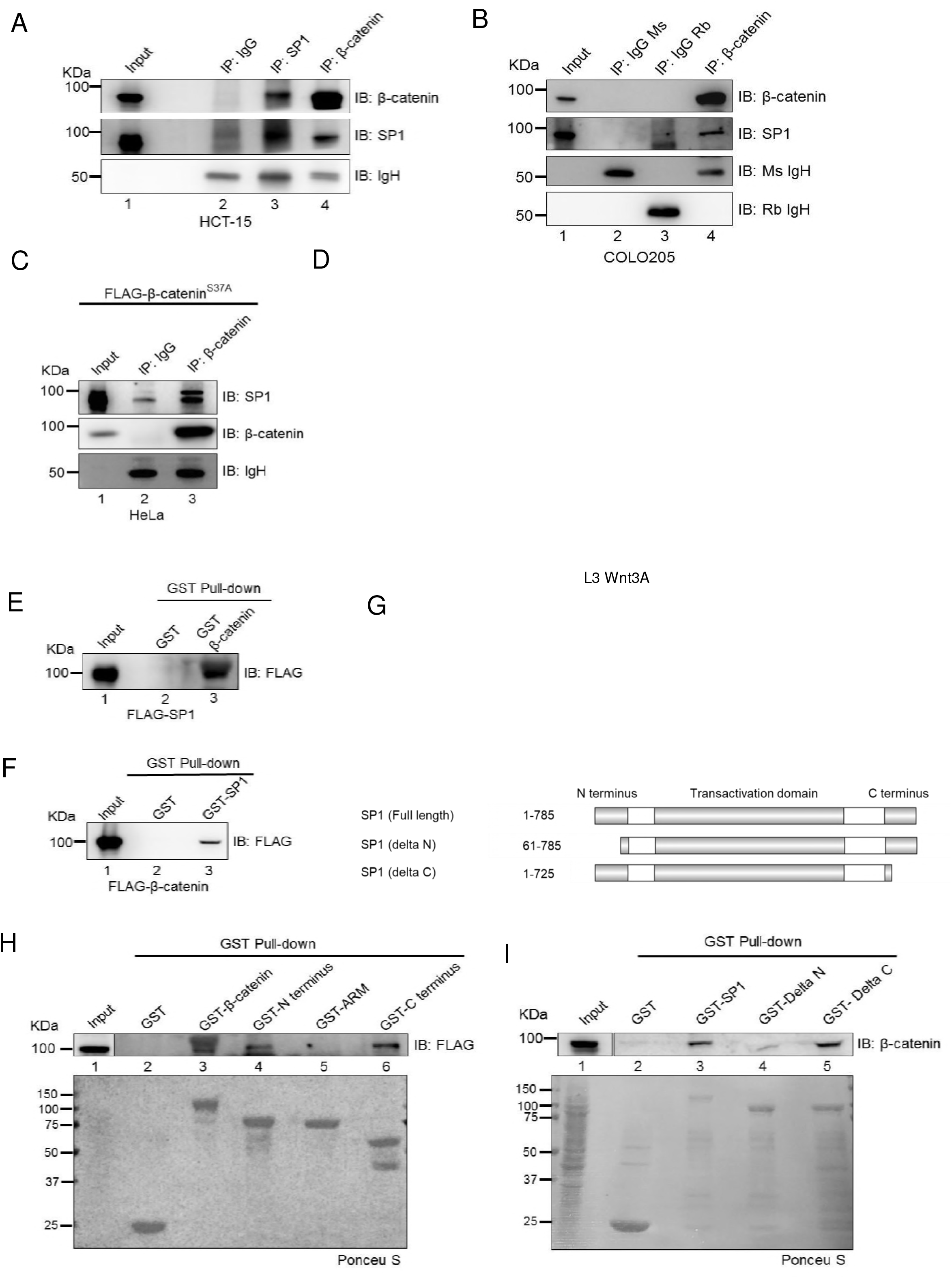
SP1 interacts with β-catenin in colorectal cancer cells. **(A)** Immunoblot for endogenous SP1 and β-catenin co-immunoprecipitated with β-catenin and SP1 respectively from HCT-15 lysates. Immunoprecipitation (IP) was done using antibody against β-catenin and SP1. IP with IgG was used as negative control. **(B)** Immunoblot for endogenous SP1 co-immunoprecipitated with endogenous β-catenin from COLO205 cells. IP with respective IgG Isotype was used as negative control. **(C)** Immunoblot for endogenous SP1 co-immunoprecipitated with β-catenin from FLAG-S37A β-catenin expressing HeLa lysate. FLAG-S37A β-catenin was overexpressed in HeLa cells and IP was done with β-catenin. IP with specific IgG Isotype used as negative control. **(D)** Representative confocal image showing colocalization of SP1 and β-catenin using SP1 and β-catenin antibodies for immunofluorescence assay in L3 Wnt3A cells. **(E)** Immunoblot for FLAG from HEK293T FLAG-SP1 expressing cells pull-down by GST β-catenin immobilized on glutathione resin. GST protein used as negative control. **(F)** Immunoblot for FLAG from HEK293T FLAG-β-catenin expressing cells pull-down by GST SP1 immobilized on glutathione resin. GST protein used as negative control. **(G)** Schematic depicting full-length and deletion constructs of β-catenin and SP1 used for in-vitro pulldown assay. Amino acid residues included in each deletion is indicated next to each construct. **(H)** Immunoblot for FLAG from HEK293T FLAG-SP1 expressing cells pulldown by GST full length β-catenin, GST N-terminus, GST ARM repeats and GST C-terminus immobilized on glutathione resin. GST protein used as negative control. **(I)** Immunoblot for FLAG from HEK293T FLAGβ-catenin expressing cells pull-down by GST full length SP1, GST delta N-terminus, and GST delta C-terminus immobilized on glutathione resin. GST protein used as negative control.

### Stabilization of β-catenin is required for Wnt signaling induced stabilization of SP1

The co-immunoprecipitation assay determined that SP1 interacts with β-catenin in Wnt signaling driven colorectal cancer cells (Figure 2A and 2B), this prompted us to investigate whether β-catenin is required for SP1 stabilization. To test this, we analyzed the expression of SP1 under β-catenin depletion in β-catenin mutant HCT116 cells. Depletion of β-catenin resulted in drastic decrease in SP1 expression at protein level (Figure 3A), but no change was observed at transcript level (Figure 3B, Supplementary Figure S3A). Immunofluorescence assay in HCT116 cells produced similar results (Figure 3C). Quantification of nuclear intensities revealed significant reduction in SP1 levels upon β-catenin depletion (Figure 3D). Furthermore, overexpression of β-catenin in SW480 cells resulted in upregulation of SP1 at protein level (Figure 3E) but not at transcript level (Figure 3F, Supplementary Figure S3B) suggesting that β-catenin is required for stability of SP1 at protein level. Next, to determine whether Wnt signaling induced SP1 stability requires β-catenin, we stimulated the FLAG-SP1 expressing HEK293 cells with Wnt3A and depleted β-catenin. Activation of Wnt signaling by Wnt3A treatment induced the stability of ectopically expressed FLAG tagged SP1 (Figure 3G). Depletion of β-catenin in FLAG-SP1 expressing cells drastically reduced the stability of SP1 even after stimulation by Wnt3A (Figure 3G, compare lanes 2 and 3). Collectively, these results suggested that β-catenin is required for Wnt signaling induced SP1 stabilization. To further corroborate the role of β-catenin in stabilization of SP1, we overexpressed FLAG-SP1 in siLuciferase HCT116 and siβ-catenin HCT116 cells. Depletion of β-catenin in FLAG-SP1 expressing HCT116 cells drastically reduced the levels of ectopically expressed FLAG-SP1 in comparison with siLuciferase FLAG-SP1 expressing cells (Figure 3H, compare lanes 2 and 3, Supplementary Figure S3C). GFP was used as transfection control. Conversely, to investigate whether β-catenin stabilization is sufficient to induce SP1 stability, we overexpressed FLAG-SP1 in control HEK293 cells and in HA-S37A β-catenin expressing cells. The phosphorylation-defective S37A mutant β-catenin is non-responsive to GSK3β mediated regulation and therefore constitutively active. The overexpression of β-catenin robustly induced the SP1 stability in comparison with vector control FLAG-SP1 expressing cells (Figure 3I, compare lanes 2 and 3, Supplementary Figure S3D). Thus, β-catenin stabilization is sufficient to induce SP1 stability.

**Figure 3:**
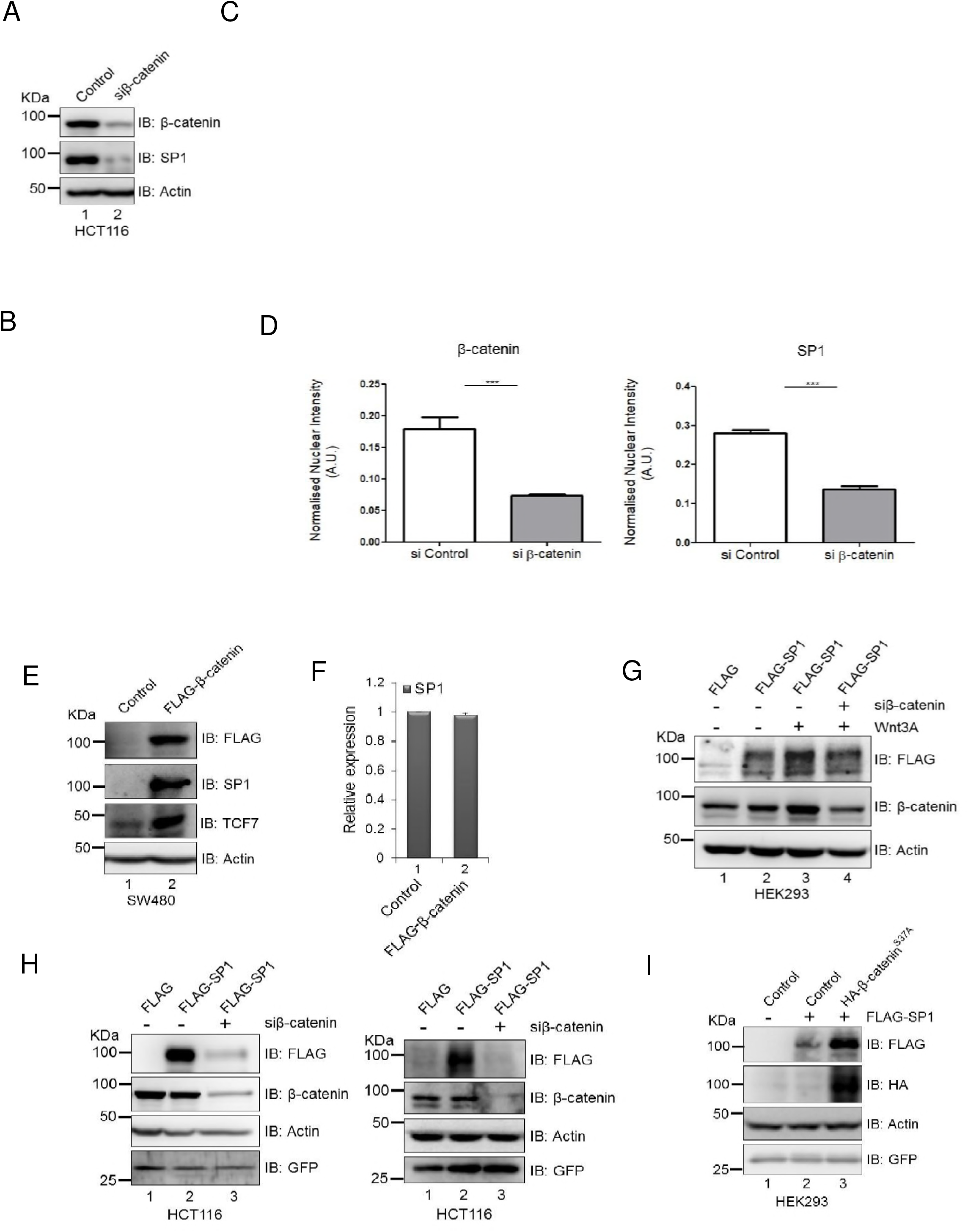
Expression of β-catenin is required for Wnt signaling induced stabilization of SP1. **(A)** Immunoblot for expression of β-catenin and SP1 in siLuciferase (control) and siβ-catenin HCT116 cells. **(B)** Relative transcript level of SP1 in sicontrol and siβ-catenin. GAPDH was used as endogenous control. Error bar represents SD for triplicates. **(C)** Immunofluorescence analysis for expression of β-catenin and SP1 in siControl and siβ-catenin transfected HCT116 cells. (Scale bar represents 20 Mm) **(D)** Quantification of nuclear intensities of SP1 and β-catenin (Mann-Whitney test, two tailed, P<0.0001). **(E)** Immunoblot for expression of SP1, TCF7 and FLAG β-catenin in control and FLAG β-catenin SW480 cells. **(F)** Relative transcript level of SP1 in control and FLAG β-catenin SW480 cells. GAPDH was used as endogenous control. Error bar represents SD for triplicates. **(G)** Immunoblot for FLAG and β-catenin in FLAG-SP1 siLuciferase and FLAG-SP1 Wnt3A treated and siβ-catenin FLAG-SP1 Wnt3A treated HEK293 cells. **(H)** Immunoblot for FLAG and β-catenin in siLuciferase, FLAG-SP1 and siβ-catenin FLAG-SP1 HCT116 cells. GFP used as transfection control (Immunoblots represent two biological replicates). **(I)** Immunoblot for FLAG and HA in control and HA β-catenin^S37A^ expressing HEK293 cells. GFP was used as transfection control.

### GSK3β is required for SP1 degradation via phoshorylation of serines in its phosphodegron motif

Next we sought to determine the mechanism by which Wnt pathway induces stability of SP1. Interestingly, in silico analysis revealed that the C-terminus of SP1 harbors a canonical phosphodegron motif that is known to be recognized by GSK3β (21) (Supplementary Figure S4A). To determine whether GSK3β could be involved in degradation of SP1 in WNT OFF state, we monitored the status of SP1 in CRL1790 primary colorectal cells treated with BIO, a GSK3β inhibitor. Inhibition of GSK3β mimicked the changes observed upon activation of Wnt signaling and robustly induced the expression of SP1 and β-catenin (Figure 4A). Next, to test whether stimulation of Wnt signaling cascade in HEK293T cells can induce SP1 stabilization at protein level, we depleted GSK3β in FLAG-SP1 expressing HEK293T cells. Depletion of GSK3β increased the level of FLAG tagged SP1 (Figure 4B, compare lanes 2 and 3, (Supplementary Figure S4B, compare lanes 2 and 3), thus demonstrating that activity of GSK3β is critical in regulating the stability of SP1. GSK3β regulates the stability of proteins through phosphorylation of the phosphodegron motif DSGXXS (21). We then argued that if such mechanism contributed to SP1 stability, mutation of the two serine residues within the phosphodegron motif at C-terminus should increase the stability of SP1 even during WNT OFF state. We mutated both serine residues to alanine (SP1^S726/732A^) and also deleted the phosphodegron containing C-terminus of SP1 (SP1^ΔDSG^). Strikingly, mutation and deletion of the canonical phosphodegron motif dramatically increased the levels of ectopically expressed FLAG-SP1 in HEK293 cells phenocopying the effect of Wnt stimulation and GSK3β inhibition (Figure 4C). To further understand the importance of phosphodegron motif, we analyzed whether SP1 from lower vertebrates such as Zebrafish, Xenopus and Chicken harbors the phosphodegron motif. Multiple sequence alignment revealed that the phosphodegron motif in SP1 is mammalian specific (Supplementary Figure S4C). To investigate whether phosphodegron is critical for stabilization of SP1, we introduced phosphodegron motif in Drosophila SP1 (dSP1) and analyzed the stability of such mutant dSP1. Interestingly, the stability of mutant is reduced in comparison with wild type dSP1 (Supplementary Figure S4D). Further, because AXIN1 is negative regulator of Wnt signaling and component of the cytoplasmic destruction complex, we investigated whether AXIN1 contributes to degradation of SP1. To test this we overexpressed AXIN1 in HCT-15 cells and monitored the expression of SP1. AXIN1 overexpression reduced the levels of SP1 (Figure 4D). Further, to understand how SP1 protein levels are reduced in response to overexpression of AXIN1, we overexpressed AXIN1 in FLAG-SP1 expressing HCT116 and SW480 cells. The overexpression of AXIN1 reduced the ectopically expressed FLAG-SP1 in comparison with control HCT116 cells (Figure 4E) and control SW480 cells (Figure 4F). Since SP1 harbors a canonical phosphodegron motif and GSK3β induces the degradation of SP1 under physiological condition and is dependent on the Wnt signaling pathway for its stabilization similar to β-catenin, we investigated whether SP1 interacts with GSK3β in WNT-OFF state. We overexpressed FLAG-SP1 in HEK293T cells and performed immunoprecipitation assay using anti- FLAG antibody. Results of co-immunoprecipitation assay suggest that SP1 physically interacts with GSK3β in WNT-OFF state (Figure 4G). We then sought to determine whether SP1 interacts with AXIN1 and is part of destruction complex under WNT OFF state. Co-immunoprecipitation analysis using HEK293T cells overexpressing FLAG-SP1 revealed that SP1 physically interacts with AXIN1 in WNT-OFF state (Figure 4H, Supplementary Figure S4E). Taken together, these results suggest that SP1 is dependent on the Wnt signaling pathway for its stabilization and interacts with components of cytoplasmic destruction complex.

**Figure 4:**
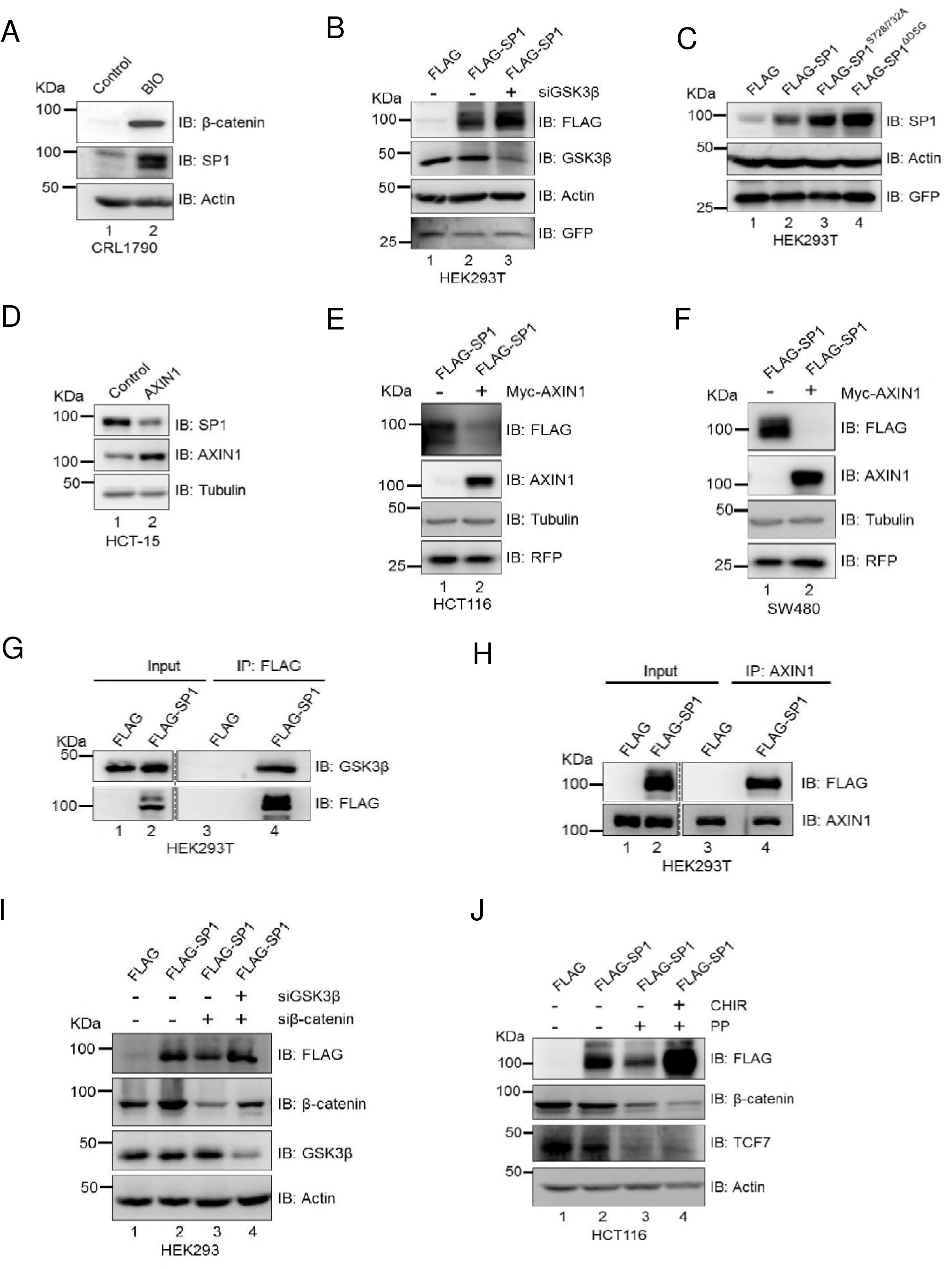
GSK3β is essential for degradation of SP1. **(A)** Immunoblot for expression of SP1 and β-catenin in control and 1 μΜ BIO, a GSK3β inhibitor and Wnt agonist. **(B)** Immunoblot for FLAG-SP1, GSK3β in siLuciferaseand in siGSK3β HEK293 cells. **(C)** Immunoblot for FLAG-SP1, FLAG-SP1 serine-alanine mutant (S726/732A) and FLAG-SP1 delta phosphodegron (ΔDSG) in HEK293 cells. GFP was used as transfection control. **(D)** Immunoblot for expression of SP1 and AXIN1 in control and Myc tagged AXIN1 expressing HCT-15 cells. Tubulin was used as endogenous control. **(E)** Immunoblot for FLAG and AXIN1 in FLAG SP1 expressing HCT116 cells upon AXIN1 overexpression. RFP was used as transfection control. **(F)** Immunoblot for FLAG-SP1 and Myc-AXIN1 in SW480 cells upon AXIN1 overexpression. RFP was used as transfection control. Tubulin was used as endogenous control in all panels. **(G)** Immunoblot for FLAG and GSK3β in FLAG-SP1 expressing HEK293T cells for co-immunoprecipitation of FLAG-SP1 with endogenous GSK3β. **(H)** Immunoblot for FLAG and AXIN1 in FLAG-SP1 expressing HEK293T cells for co-immunoprecipitation of FLAG-SP1 with endogenous AXIN1. Immunoprecipitation was performed using antibody against endogenous AXIN1. **(I)** Immunoblot analysis in FLAG-SP1 expressing HEK293 cells transfected with siβ-catenin alone (lane 3) and with siGSK3β (lane 4). Actin was used as endogenous control. **(J)** Immunoblot for FLAG, β-catenin and TCF7 in control FLAG-SP1 HCT116 cells, PP (Pyrvinium Pamoate) treated FLAG-SP1 expressing HCT116 cells and FLAG-SP1 expressing HCT116 cells treated with both PP and CHIR. Actin was used as endogenous control.

We observed that GSK3β induces SP1 degradation in WNT-OFF state and also overexpression of β-catenin reverses the effect of GSK3β by stabilization of SP1 even in WNT-OFF state. This prompted us to investigate whether depletion of β-catenin induces degradation of SP1 via GSK3β. Towards this, we monitored SP1 stability in HEK293 cells in which FLAG-SP1 was overexpressed and β-catenin was depleted or alternatively, both GSK3β and β-catenin were depleted simultaneously. Depletion of β-catenin reduced the levels of ectopically expressed FLAG-SP1 in comparison with siControl (Figure 4I, compare lanes 2 with 3). In contrast, depletion of GSK3β in β-catenin depleted FLAG-SP1 expressing HEK293 cells rescued the ectopic levels of FLAG-SP1 (Figure 4I, compare lane 3 with lane 4). Further, to understand this cross talk, we analyzed SP1 stability upon β-catenin degradation induced by Pyrvinium Pamoate (PP), a known inhibitor of Wnt signaling (22) and also inhibited GSK3β activity in PP treated FLAG-SP1 expressing cells using CHIR. PP treatment induces the degradation of β-catenin so as that of FLAG tagged SP1 (Figure 4J, compare lanes 2 and 3, Supplementary Figure S4F), whereas inhibition of GSK3β by CHIR restored the levels of FLAG-SP1 (Figure 4J, compare lanes 3 and 4). These results demonstrate that β-catenin is essential to prevent the degradation of SP1 mediated by GSK3β activity. Interestingly, HCT116 cells possess functional destruction complex, however, with a mutation in β-catenin (Serine 45) to evade the effect of destruction complex. Thus, higher levels of β-catenin are maintained in these cells, presumably explaining why SP1 levels are higher in HCT116 cells even after having active GSK3β and functional destruction complex. These findings therefore establish mechanistic link for requirement of β-catenin to maintain stabilization of SP1. Notably, the degradation of β-catenin leads to downregulation of the known Wnt-responsive gene TCF7. However, rescue of SP1 expression by CHIR treatment did not restore the expression of TCF7 (Figure 4J).

### Active Wnt signaling impedes interaction of SP1 with β-TrCP and not with GSK3β

GSK3β regulated proteins like β-catenin undergo phosphorylation of serine residues in phosphodegron motif and is recognized and ubiquitinated by F box containing protein β-TrCP ubiquitin E3 ligase followed by degradation through proteosomal degradation pathway. We therefore sought to determine whether Wnt signaling impedes the interaction of SP1 with GSK3β and β-TrCP in HEK293 cells. To test this, we treated FLAG-SP1 expressing cells with Wnt3A. Treatment with soluble Wnt3A ligand robustly stabilized FLAG-SP1 in HEK193 cells (Figure 5A, compare lanes 2 and 3). Notably, overexpression of SP1 also induced the expression of β-catenin (Figure 5A, compare lanes 1 and 2). Similarly Wnt3A treatment further increased the β-catenin expression indicating the activation of Wnt signaling (Figure 5A, compare lanes 2 and 3). Next, we immunoprecipitated β-TrCP in FLAG-SP1 expressing cells upon Wnt stimulation. The co-immunoprecipitation analysis revealed that Wnt stimulation impedes the interaction of SP1 with β-TrCP (Figure 5B, compare lane 5 and lane 6 in IP blot). To determine whether Wnt stimulation also abrogates the SP1 interaction with GSK3β, we immunoprecipitated FLAG-SP1 expressing cells with anti- FLAG antibody in control and Wnt3A treated cells and performed co-immunoprecipitation analysis. We observed that Wnt stimulation did not impede the interaction of SP1 with GSK3β (Figure 5C, compare lanes 5 and 6). Thus indicating that Wnt stimulation only prevents interaction of β-TrCP, not GSK3β with SP1, thereby evading the recognition by proteosomal pathway post ubiquitination by β-TrCP. To further corroborate the role of aberrant activation of Wnt signaling in SP1 stabilization, we inactivated GSK3β by CHIR treatment, cells were also treated with MG132 and analyzed for the interaction of GSK3β with SP1. Co-immunoprecipitation analysis revealed that although CHIR treatment results in robust increase in SP1 stabilization, it did not impede the interaction of SP1 with GSK3β (Figure 5D, compare lanes 5 and 6). Further, we thought to investigate the interaction of GSK3β with SP1 and its phosphodegron mutant. Towards this, co-immunoprecipitation was performed using lysates of HEK293T cells overexpressing wild-type FLAG-SP1 and mutant FLAG-SP1. Co-immunoprecipitation analysis suggested that mutant SP1 interacts with lesser affinity with GSK3β in comparison to the wild-type SP1 (Supplementary Figure S5, compare lanes 5 and 6). These results indicate that SP1 remains bound to GSK3β even after inactivation of GSK3β activity but loses interaction with β-TrCP. Since β-catenin induces stabilization of SP1 and mimics the stabilization induced upon Wnt stimulation, we analyzed interaction of SP1 with β-TrCP upon Wnt stimulation and mutant β-catenin overexpression. Immunoblot analysis revealed that both Wnt stimulation and β-catenin overexpression impede the interaction of SP1 with β-TrCP, thereby inducing stabilization of SP1 (Figure 5E). Wnt stimulation, via affecting the interaction of SP1 with β-TrCP, presumably prevents its ubiquitination and subsequent proteosomal degradation. To investigate this, we analyzed the *in vivo* ubiquitination of SP1 upon Wnt stimulation. We overexpressed FLAG-SP1 in HA-ubiquitin expressing HEK293 control cells and Wnt3A treated cells. Co-immunoprecipitation analysis revealed loss of SP1 ubiquitination upon Wnt stimulation (Figure 5G). To extend the role of Wnt signaling in stabilization of SP1, we analyzed the interaction SP1 with AXIN1 upon Wnt stimulation. The data reveals that Wnt stimulation impedes the interaction of SP1 with AXIN1 (Figure 5F, compare lanes 5 and 6). These results provide a novel link that in WNT-ON cells, Wnt stimulation prevents interaction of AXIN1 and β-TrCP with SP1 thereby preventing ubiquitination and inducing its stability. In contrast, under WNT-OFF state GSK3β mediated phosphorylation of serine residues within the phosphodegron motif allows recognition of SP1 by β-TrCP and subsequent degradation by proteosomal pathway.

**Figure 5:**
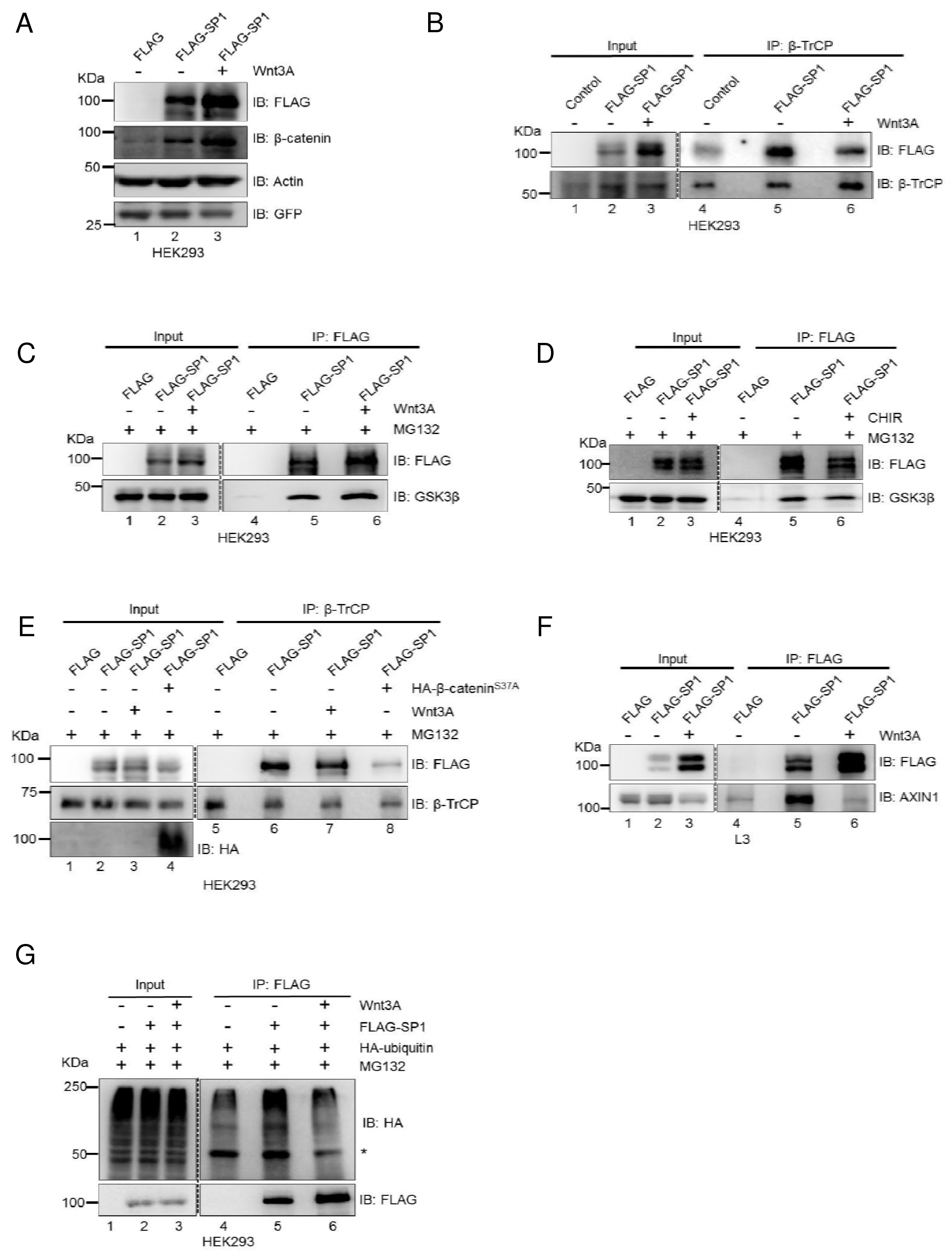
Wnt signaling stimulation abrogates SP1’s interaction with β-TrCP and not with GSK3β. **(A)** Immunoblot using anti- FLAG antibody in FLAG control, FLAG-SP1 and FLAG-SP1 Wnt3A treated HEK293 cells. HEK293 cells were treated with Wnt3A for 6h. Actin used as endogenous control. GFP used as transfection control. **(B)** Immunoblot for FLAG and β-TrCP for co-immunoprecipitation of FLAG-SP1 with endogenous β-TrCP from FLAG, FLAG-SP1 and FLAG-SP1 Wnt3A treated HEK293 cells. IP with β-TrCP was performed in HEK293 cells overexpressing FLAG, FLAG-SP1 control and FLAG-SP1 + Wnt3A treatment. **(C)** Immunoblot for endogenous GSK3β co-immunoprecipitation with FLAG-SP1 from FLAG and FLAG-SP1 HEK293 cells. IP was performed using anti- FLAG antibody. HEK293 cells were treated with 10 μM MG132 along with Wnt3A for 6 h. **(D)** Immunoblot for endogenous GSK3β co-immunoprecipitation with FLAG-SP1 from FLAG, FLAG-SP1, and FLAG-SP1 CHIR treated HEK293 cells. IP was performed using antiFLAG antibody. HEK293 cells were treated with 10 μM MG132 along with 3 μM CHIR for 6h. **(E)** Immunoblot for FLAG, β-TrCP and HA for co-immunoprecipitation of FLAG-SP1 with endogenous β-TrCP in FLAG, FLAG-SP1 control, FLAG-SP1 Wnt3A treated HEK293 cells and FLAG-SP1 + S37A β-catenin overexpressing HEK293 cells. IP was performed using anti- β-TrCP antibody. **(F)** Immunoblot for FLAG and AXIN1 for co-immunoprecipitation of FLAG-SP1 with endogenous AXIN1 in FLAG, FLAG-SP1 control and FLAG-SP1 Wnt3A conditioned media treated L3 cells. **(G)** Immunoblot for HA and FLAG for co-immunoprecipitation of FLAG SP1 with HA-ubiquitin in HA-ubiquitin expressing FLAG, FLAG-SP1 control and FLAG-SP1 Wnt3A treated HEK293 cells. ^*^ IgG heavy chain.

### SP1 is required for Wnt signaling-dependent β-catenin stabilization and regulation of Wnt responsive genes

The common pathway regulating SP1 and β-catenin suggested and immunofluorescence analysis in HCT116 cells revealed significant reduction in β-catenin levels upon SP1 depletion (Figure 6A). Quantification of nuclear intensities revealed significant reduction in β-catenin levels upon SP1 depletion (Figure 6B). Similarly, overexpression of SP1 in SW480 cells induced β-catenin expression (Supplementary Figure S6A). These observations prompted us to investigate whether SP1 is required to stabilize β-catenin. To investigate this, we treated siControl HEK293 cells and siSP1 HEK293 cells with Wnt3A for 6h. Wnt stimulation induced the stability of both SP1 and β-catenin (Figure 6C, compare lane 1 with lane 2), whereas SP1 depletion in Wnt stimulated HEK293 cells was sufficient to reduce the levels of β-catenin so as the expression of the known Wnt responsive gene TCF7 (Figure 6C, compare lanes 2 and 3). Further, CHIR induced expression of β-catenin in CRL1790 was reduced upon depletion of SP1 in comparison with control (Figure 6D, compare lanes 2 and 3). To test whether SP1 is required to stabilize β-catenin at protein level, we investigated the stability of ectopically expressed FLAG β-catenin in HeLa cells upon SP1 depletion. We overexpressed FLAG β-catenin in siLuciferase and siSP1 cells. Overexpression of β-catenin resulted in robust increase in SP1 expression (Figure 6E, compare lanes 1 and 2), whereas depletion of SP1 reduced the levels of ectopically expressed FLAG β-catenin (Figure 6E, compare lanes 2 and 3 anti-FLAG blot). Similar observations were recorded in HEK293 cells wherein SP1 depletion markedly reduced the expression of ectopically expressed FLAG β-catenin (Figure 6F compare lanes 2 and 3; Supplementary Figure S6B, compare lanes 2 and 3, Supplementary Figure S6C and S6D). Next, to investigate whether SP1 is sufficient to stabilize β-catenin, we overexpressed Myc tagged β-catenin in control and phosphodegron mutant SP1 expressing HEK293 cells. The overexpression of mutant SP1 induced the levels of ectopically expressed Mycβ-catenin in comparison with control (Figure 6G, compare lanes 2, 3 and 4). Surprisingly, overexpression of mutant SP1 also promoted the stabilization of S37A mutant β-catenin, indicating GSK3 independent role of SP1 in β-catenin stabilization (Supplementary Figure S6E). The requirement of SP1 for β-catenin stabilization prompted us to investigate whether SP1 abolishes the interaction of β-catenin with components of destruction complex and thereby β-TrCP mediated ubiquitination. To investigate this hypothesis, we overexpressed mutant SP1 in HA ubiquitin expressing HEK293T cells and performed *in vivo* ubiquitination followed by co-immunoprecipitation assays. Overexpression of mutant SP1 abolished the β-catenin interaction with AXIN1, GSK3β and β-TrCP (Figure 6H, compare lane 5 and lane 6) and abolished β-TrCP mediated ubiquitination (Figure 6H, compare lanes 5 and 6). Additionally, co-immunoprecipitation data established that stabilized SP1 impedes the recruitment of GSK3β to AXIN1 thereby inducing the destabilization of the destruction complex (Supplementary Figure S6F, compare lanes 5 and 6). To further understand if SP1 is required for β-catenin stabilization, we generated a stable β-catenin overexpressing HEK293 cell line and induced stabilization of β-catenin by treating with CHIR. Inactivation of GSK3β induced the stability of FLAGβ-catenin as well as that of SP1 (Figure 6I, compare lanes 1 and 2), whereas SP1 depletion in FLAG β-catenin expressing cells reduced the levels of ectopically expressed FLAG β-catenin (Figure 6I compare lanes 2 and 3). The activation of Wnt signaling resulted in upregulation of the Wnt responsive genes AXIN2 and TCF7 whereas SP1 depletion was sufficient to reduce their expression significantly. To test whether SP1 depletion mediated degradation of FLAG tagged β-catenin at protein level occurs via proteosomal pathway, we treated CHIR treated FLAGβ-catenin SP1 depleted cells with MG132 (Fig. 6I, lane 4). MG132 treatment rescued the stabilization of FLAGβ-catenin (Figure 6I, compare lanes 3 and 4). Interestingly, the rescue of β-catenin stabilization did not re-induce the expression of Wnt responsive genes, suggesting the role of SP1 in regulation of Wnt responsive genes. This was further confirmed by the Wnt specific TOP/FOP reporter activity in CHIR treated SP1 depleted cells. CHIR treatment induced the reporter activity (Figure 6J, compare lanes 1 and 2). The depletion of SP1 in CHIR treated cells was sufficient to reduce the TOP/FOP reporter activity significantly (Figure 6J, compare lanes 2 and 3). To further establish the dual role of SP1 in β-catenin stabilization and regulation of Wnt responsive genes, we overexpressed mutant β-catenin in dose-dependent manner in siSP1 HEK293 cells to overcome the effect of SP1-dependent β-catenin stabilization. Overexpression of β-catenin induced the stability of SP1 as well as the expression of Wnt responsive genes AXIN2 and TCF7, whereas the depletion of SP1 reduced the levels of Wnt responsive genes even after rescuing the levels of β-catenin (Figure 6K). TCF7L2 is a major transcription factor driving the expression of Wnt responsive genes (23). Lack of any noticeable change in TCF7L2 expression in SP1 depleted cells rules out the possibility that decrease in expression of Wnt responsive genes upon SP1 depletion could be through regulation of TCF7L2 (Figure 6K).

**Figure 6:**
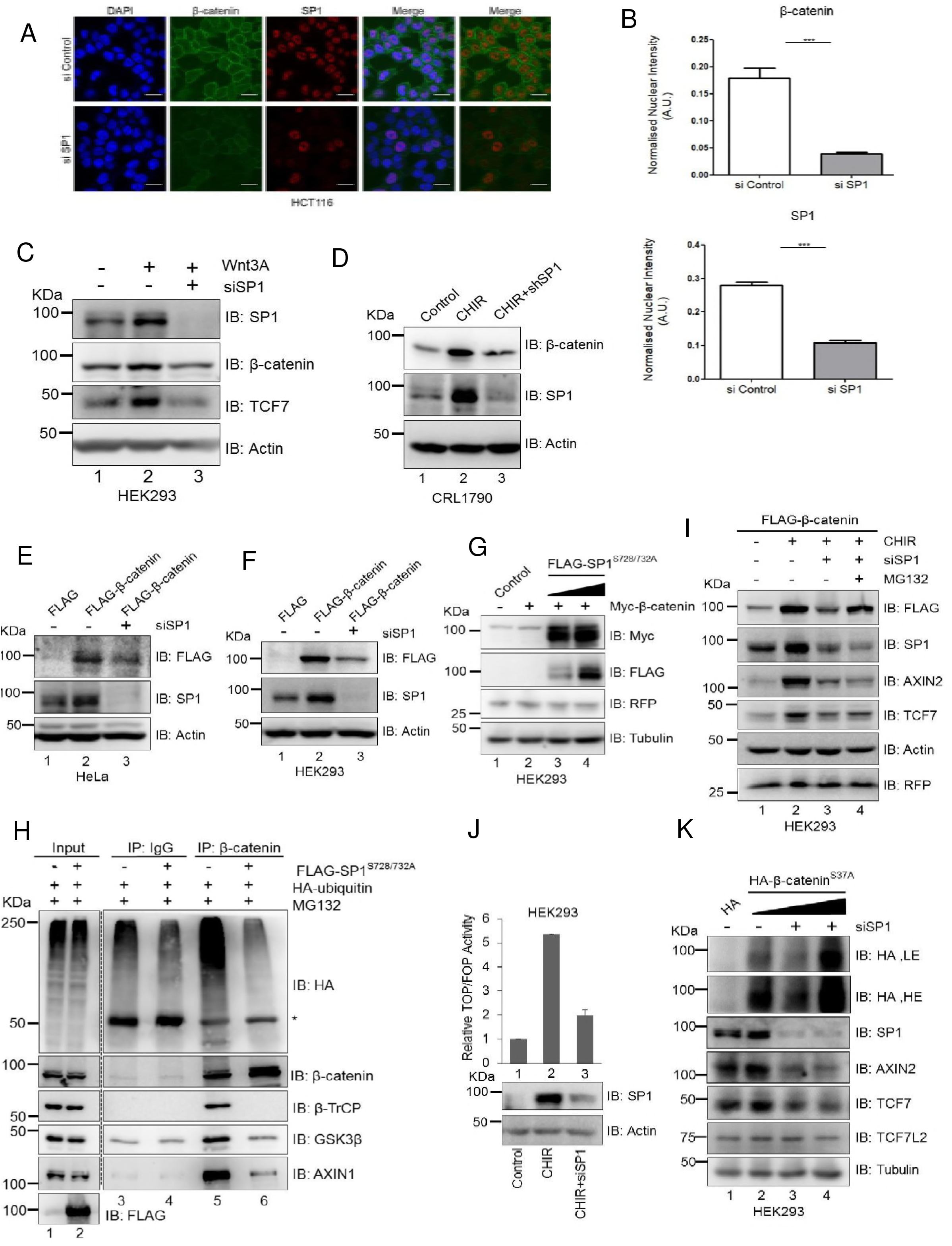
SP1 is required for Wnt signaling-dependent β-catenin stabilization and regulation of Wnt responsive genes. **(A)** Immunofluorescence analysis for expression of β-catenin and SP1 in sicontrol and siSP1 transfected HCT116 cells. (Scale bar represents 20 μm) **(B)** Quantification of nuclear intensities of SP1 and β-catenin (Mann-Whitney test, two tailed, P<0.0001) **(C)** Immunoblot for SP1 and β-catenin upon Wnt3A treatment in control and SP1 depleted HEK293 cells. HEK293 cells were transfected with siGFP and siSP1, treated with Wnt3A after 42 h for 6 h. **(D)** Immunoblot for SP1 and β-catenin upon CHIR treatment in control and SP1 depleted CRL-1790 cells. CRL-1790 cells were transfected with siGFP and siSP1, treated with CHIR for 48 h **(E)** Immunoblot for FLAG and for SP1 in FLAGβ-catenin si luciferase HeLa cells and FLAG-β-catenin siSP1 cells. **(F)** Immunoblot for FLAG and for SP1 in FLAGβ-catenin siLuciferase HEK293T cells and FLAGβ-catenin siSP1 cells **(G)** Immunoblot for MYC and FLAG using FLAG in MYC-β-catenin control and MYC-β-catenin FLAG-SP1 mutant HEK293 cells. RFP was used for transfection control. **(H)** Immunoblot for HA, AXIN1, GSK3β, β-TrCP and β-catenin in control HA-ubiquitin and HA-Ubiquitin FLAG mutant SP1 HEK293 for co-immunoprecipitation of β-catenin with ubiquitin, AXIN1, GSK3β and β-TrCP. Mouse IgG as negative control. ^*^ IgG heavy chain. **(I)** Immunoblot for FLAG, AXIN2, TCF7 and SP1. Stable FLAGβ-catenin HEK293 cells were treated with CHIR in sicontrol, siSP1 and siSP1 MG132 HEK293 cells. RFP used as transfection control. **(J)** Graphical representation of TOP/FOP reporter activity under CHIR treatment in sicontrol and siSP1 HEK293 cells. Below graph, Immunoblot for reporter assay. **(K)** Immunoblot for HA, SP1, AXIN2 and TCF7 in control and in SP1 depleted HEK293 cells upon S37A β-catenin overexpression in dose-dependent manner. Tubulin was used as endogenous control for all immunoblots.

### SP1 binds to promoters of TCF7L2/β-catenin target genes

To elucidate how stability of SP1 could be involved in regulating Wnt responsive genes, we analyzed the ChIP-seq data for genome-wide occupancy of SP1 in HCT116 colorectal cancer cells from the ENCODE consortium. We identified SP1 binding peaks on promoters of TCF7L2/β-catenin target genes AXIN2, c-Myc and TCF7 (Figure 7A, 7B and 7C, Supplementary Figure S7 A and B). To directly demonstrate role of SP1/β-catenin signaling in regulation of Wnt responsive genes, we performed chromatin immunoprecipitation (ChIP) in HCT116 cells using SP1 and β-catenin antibodies and analyzed the expression of key Wnt responsive genes AXIN2 and TCF7 at transcript level. ChIP analysis demonstrated the co-occupancy of SP1 and β-catenin on promoters of AXIN2 and TCF7 (Figure 7D). Quantitative gene expression profiling revealed that SP1 depletion robustly reduced the expression of Wnt responsive genes (Figure 7E). These findings suggest that SP1 and β-catenin presumably co-occupy the promoters of Wnt responsive genes and regulate their expression. These results further establish direct mechanistic role of SP1 in regulating β-catenin stabilization and regulation of Wnt responsive genes in β-catenin-dependent manner. Collectively, these results strongly argue for a mechanism wherein SP1 and β-catenin are required for mutual stabilization in Wnt signaling-dependent manner and for regulation of Wnt responsive genes (Figure 7F).

**Figure 7:**
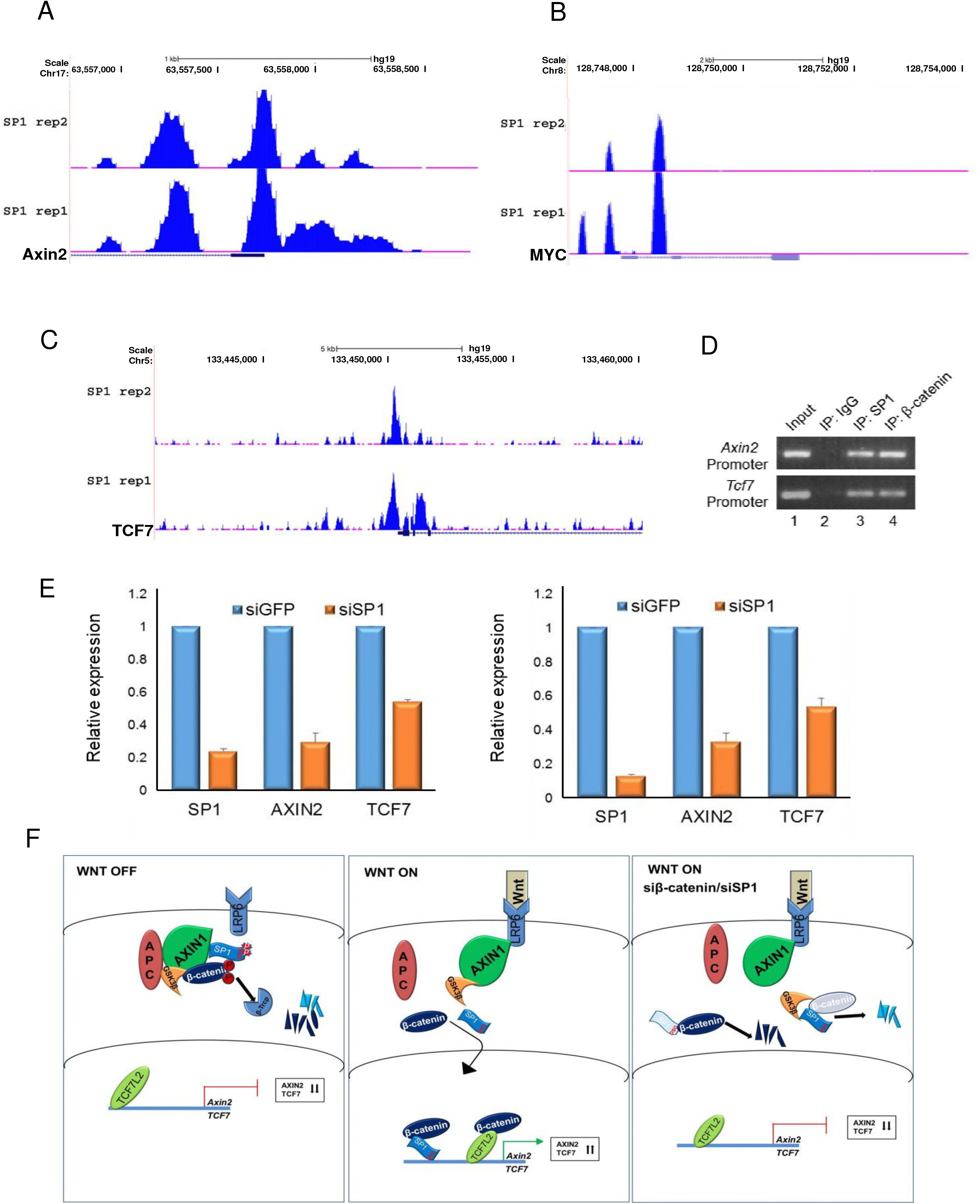
SP1/β-catenin directly bind to the promoters of Wnt responsive genes. **(A)** UCSC genome browser view showing occupancy of SP1 for two independent replicates on *Axin2, c-Myc* **(B)** and *Tcf7* promoters **(C)**. ChIP-seq data was analyzed as described in ‘Methods’. **(D)** Chromatin immunoprecipitation assay showing co-occupancy of SP1 and β-catenin on promoters of AXIN2 and TCF7. **(E)** Quantitative RT-PCR analysis to monitor expression of Wnt responsive genes upon SP1 depletion in HCT-116 cells. The two graphs represent data from two biological replicates. Error bars represent standard deviation for technical replicates. **(F)** Model depicting Wnt signaling mediated stabilization of SP1, and SP1 and β-catenin mutual stabilization. In WNT-OFF state, SP1 and β-catenin are degraded in GSK3β-dependent manner. Decreased expression of SP1 and β-catenin is reflected in reduced expression of Wnt responsive genes. In WNT-ON state, both SP1 and β-catenin are stabilized. Increased expression of SP1 and β-catenin induce the expression of Wnt responsive genes. In WNT-ON state, depletion of β-catenin induces the degradation of SP1 and depletion of SP1 induces the degradation of β-catenin, thereby reversing the Wnt induced cellular phenotype of increased expression of Wnt responsive genes.

## Discussion

Here, we present a novel mode of regulation of Wnt signaling involving crosstalk of SP1 and β-catenin stabilization as integral part of Wnt signaling cascade. We show that Wnt stimulation induces the simultaneous stabilization of SP1 and β-catenin. Further, using various biochemical inhibitors of downstream effectors we demonstrate that Wnt signaling induces SP1 stabilization. After demonstrating that SP1 and β-catenin follow similar events upon activation of canonical Wnt signaling, we investigated whether SP1 is component of the β-catenin destruction complex. We show that SP1 interacts with AXIN1 and GSK3β in Wnt-OFF state. Further, in colorectal cancer cells characterized by constitutively active Wnt signaling, SP1 interacts with β-catenin and is part of the destruction complex.

### β-catenin promotes Wnt signaling-dependent stabilization of SP1

Here we provide evidence towards a novel regulatory complex in Wnt signaling wherein not only β-catenin is stabilized but transcription factor SP1 also follows the same route to stabilization. This study sheds light on the hitherto uncharacterized role of Wnt signaling in stabilization of SP1. We show that in absence of Wnt ligand (WNT OFF state) SP1 interacts with components of the destruction complex, suggesting destruction complex is critical for SP1 destabilization. Stimulation by Wnt ligand (WNT ON state) leads to loss of interaction of SP1 with β-TrCP and AXIN1 but not with GSK3β leading to increase in levels of SP1 in cytosol and nucleus. This change in SP1 stabilization is similar to β-catenin stabilization whereby Wnt stimulation impedes interaction of β-catenin with β-TrCP but not with GSK3β (24).

After understanding the importance of Wnt signaling and destruction complex, we focused on understanding the role of β-catenin in stabilization of SP1. Inhibition of cytoplasmic destruction complex leads to stabilization and nuclear translocation of β-catenin. Here we report role of β-catenin in WNT OFF and WNT ON state. We provide evidences that in WNT ON state stabilization of β-catenin results in increase in SP1 at protein level but transcript levels are unaffected. We show that stabilization of SP1 is dependent on that of β-catenin. In WNT OFF state the stabilization of β-catenin mimics the changes acquired upon Wnt stimulation by impeding the interaction of SP1 with the E3 ubiquitin ligase β-TrCP. These results suggest that both SP1 and β-catenin form complex and release of β-catenin from destruction complex upon Wnt stimulation prevents recruitment of SP1 to the destruction complex. The dependence of SP1 stabilization on β-catenin could be very critical for therapeutic intervention in cancers driven by SP1 hyperexpression. Small molecules that are used to target Wnt signaling driven cancers can presumably be used to target SP1 driven cancers. Furthermore, SP1 has been shown to be associated with cancers driven by aberrant activation of Wnt signaling (25–27). Thus, it is of immediate interest to understand how this novel cross talk of SP1 and β-catenin could be critical for targeted drug therapy in cancers driven by SP1/β-catenin signaling. It would be important to elucidate whether SP1 can modulate the function of novel players such as YAP and SMADs that are recently shown to exhibit crosstalk with Wnt signaling (28, 30).

### GSK3β mediated SP1 degradation and role of phosphodegron motif

GSK3β is critical for regulating stabilization of proteins containing the DSGXXS motif also referred to as the phosphodegron motif (2, 23, 28) Phosphorylation of serine residues in this motif is recognition tag for β-TrCP to ubiquitinate, followed by proteosomal degradation (7). We observed that GSK3β mediated destabilization of SP1 requires loss of β-catenin. We provide evidences that SP1 interacts with both GSK3β and β-catenin. Further, stabilization of β-catenin impedes interaction of SP1 with components of the destruction complex and destabilization of SP1 upon β-catenin depletion can be rescued by depleting or inhibiting GSK3β. Thus, availability of β-catenin is essential for SP1 stabilization and GSK3β can promote SP1 degradation only in absence of β-catenin. Further, in WNT OFF state, availability of GSK3β is critical to induce SP1 degradation and depletion of GSK3β mimics the changes observed upon stabilization of β-catenin and/or upon Wnt stimulation. The degradation of SP1 upon β-catenin depletion can be reversed by depleting GSK3β or inactivating its kinase activity. We propose that in the absence of β-catenin or in the Wnt-OFF state, GSK3β engages SP1 for phosphorylation of its phosphodegron motif which is subsequently ubiquitinated by E3 ubiquitin ligase followed by proteasome mediated degradation. The importance of phosphodegron in SP1 is also underscored by fact that it is specific only to mammals. The introduction of mammalian phosphodegron in Drosophila SP1 resulted in its destabilization.

Earlier studies have shown that Thiazolidinedione drugs induce expression of β-TrCP thereby destabilize SP1 in prostate cancers (27). Further, Thiazolidinediones has been shown to induce degradation of β-catenin in GSK3β-dependent manner (31), thus hinting at the common route to stabilization. Here we demonstrate a connecting link that co-regulates SP1 and β-catenin. We report that β-catenin stability is essential for SP1 stability and function of GSK3β is critical for this regulation. Further, we show that N- and C-termini of β-catenin interact with N-terminus of SP1 leaving C-terminus containing phosphodegron motif available for interaction with GSK3β. Upon Wnt stimulation, SP1 fails to interact with GSK3β. These results therefore indicate that availability of β-catenin functions as a strong molecular cue regulating SP1 stabilization. Further investigation is required to delineate the molecular mechanism governing recruitment of β-TrCP to SP1. A recent study revealed that YAP is required for recruitment of β-TrCP to β-catenin in WNT OFF state, hence inducing β-catenin degradation (28). It will be interesting to test whether YAP is a common mediator for β-catenin and SP1 regulation in WNT OFF state.

### SP1 promotes β-catenin stabilization

We further established that SP1 could mimic Wnt signaling associated stabilization of β-catenin. We provide evidences that Wnt/GSKβ signaling induced stabilization of β-catenin is attenuated in absence of SP1. The common route of SP1 and β-catenin stabilization by Wnt stimulation seems to be critical for their mutual stabilization. Further, we found that by overexpressing stabilized phosphodegron mutant SP1 in WNT OFF state leads to stabilization of β-catenin by impeding its recruitment to the cytoplasmic destruction complex. The stabilization of SP1 seems to be critical for regulating stabilization of destruction complex such that it impedes the interaction of GSK3β to AXIN1. The requirement of SP1 in the Wnt signaling cascade might be critical for stringent regulation during embryonic development and progression of Wnt driven cancers. Hence understanding the molecular mechanism of how SP1 regulates interaction of β-catenin with components of the destruction complex will be useful for targeting drugs against developmental defects and cancers. The SP1 and β-catenin are required for reciprocal stabilization where stabilization of one mediates the changes required to induce stabilization of other, thereby exerting regulatory feedback (See schematic model representation in Figure 7F). The mutual sequestration targeting SP1 and β-catenin complex could be important for therapeutic intervention in future. Understanding the interaction/crosstalk of SP1/β-catenin signaling axis with other signaling pathways could provide further mechanistic insights into tumorigenesis.

### Stability of SP1 affects the outcome of Wnt signaling

We provide evidence that SP1 is not only required to stabilize β-catenin but is also essential to regulate the expression of Wnt responsive genes. There are two changes associated with SP1 expression. In first scenario, SP1 interacts with β-catenin and is required for its Wnt dependent regulation of stabilization. In second scenario, nuclear role of SP1 is critical wherein it co-occupies with β-catenin the promoters of Wnt responsive genes and regulates their expression. The importance of SP1 expression can be attributed by the fact that even rescuing the levels of β-catenin in SP1 depleted cells does not re-induce the expression of Wnt responsive genes. These molecular changes that govern the expression of SP1/β-catenin signaling-dependent regulation of Wnt responsive genes are independent of TCF7L2 expression. Taken together, these findings suggest that SP1 acts as a double-edged sword that is required for stabilization of β-catenin and for regulation of Wnt responsive genes and therefore is critical determinant of molecular changes associated with Wnt signaling pathway.

SP1 induces transcriptional super-activation by tetramerization through its D domain (32–34). This mode of SP1 organization mediates synergistic activation or repression of genes. Further, tetrameric aggregation of SP1 has been shown to induce DNA looping between enhancer and promoter regions of genes (35). The tetrameric assembly of SP1 on DNA provides docking site for other transcription factors, transcriptional regulators and chromatin remodelers (24), thus providing an essential platform to activate or repress gene expression. Hence it will interesting to understand whether interaction of SP1 with β-catenin influences such arrangement on promoters of Wnt responsive genes along with TCF7L2/β-catenin complex to super-activate the expression of target genes upon Wnt stimulation. Further, it will be interesting to understand whether differential gene expression regulated by SP1 depends on β-catenin co-occupancy on target genes. Genome-wide approaches would reveal the differential role of SP1 and SP1/β-catenin complex for regulation of gene expression and thereby influencing the different developmental paradigms and progression of cancers.

## Materials and Methods

### Cell culture, transfections and western blotting

SW480, SW620 and HCT116 cell lines were grown in DMEM with 10% FCS. CRL1790 were grown in MEM with 10% FCS. CRL1790, SW480, SW620 and HeLa cell lines were obtained from American Type Culture Collection (ATCC). HCT116 was obtained from European Collection of Cell Cultures (ECACC through SIGMA). For transfection with siRNA mediated depletion, cells were seeded and transfected after 24 h with indicated siRNA and then harvested for immunoblotting and RNA extraction after another 48 h. The sequences of shRNA and siRNA used are listed in supplementary Table 1.

### Treatments with inhibitors

To activate Wnt signaling CRL1790 cells were treated with CHIR (3 μM) and BIO (1 μM) for 48 h and harvested for protein and RNA. For stability experiments cells were first transfected with siRNA and after 24 h transfected with plasmids and/or treated with Wnt3A (100 ng/ml) or (CHIR 3 μM/ml) as mentioned in figure legends. Similarly, Pyrvinium Pamoate (PP) was used at concentration of 100 nM and cells were harvested after 48 h. For knockdown of SP1 and β-catenin under CHIR and Wnt3A, cells were first transfected with control siRNA, siSP1 and siβ-catenin and after 12 h treated with CHIR for 48 h in case of CRL1790 cells, whereas other cells were treated with CHIR and Wnt3a for 6 h after 42 h of transfection. ALLN was used at 25 μg/ml. MG132 was used at 6 μM/ml.

### Immunofluorescence analysis and quantification

Cells were cultured on Fibronectin coated glass coverslips and fixed using 4% paraformaldehyde (PFA) in PBS for 15 min at room temperature (RT). Permeabilization was performed using 0.25% Triton X-100 for 5 minutes, followed by blocking in 10% BSA in PBS for 1 h at RT. The cells were incubated in primary antibody (diluted in 5% BSA in PBS) for 16 h at 4 °C followed by four washes in PBS at RT, incubation in secondary antibody (diluted in 2% BSA) for 1 h at RT and four washes in PBS at RT. Confocal images were obtained using Zeiss 710 LSM confocal microscope. Nuclear intensity was quantified using ImageJ software. Nuclear area within each cell was selected using DAPI staining as the region of interest (ROI). Quantified nuclear intensity was normalized with the nuclear area and DAPI. At least 300 cells were used for quantification in each set.

### ChIP-seq data analysis

Genome-wide occupancy datasets of SP1 in HCT116 cells were obtained from publicly available database from ENCODE consortium with experiment IDs ENCSR000BVT and ENCSR000BSF for Input and SP1 ChIP samples respectively. Briefly, reads were aligned to human genome (hg19) with bowtie2 using very-sensitive mode. Peaks were called using MACS14 and wiggle files were uploaded on the UCSC genome browser for visualization.

### Antibodies, reagents and plasmids

SATB1, TCF7L2, TCF7, SP1, GSK3β, β-TrCP, AXIN1 and AXIN2 antibodies were obtained from Cell Signaling Technology. Anti- β-catenin antibody was obtained from BD Transduction Laboratories. β-catenin for IP in HeLa was used from Santa Cruz. Actin and gamma-tubulin antibodies were obtained from Sigma. Secondary HRP conjugated antibodies were obtained from BioRad. Recombinant Wnt3A protein was obtained from R&D systems. Pyrvinium Pamoate was obtained from Sigma. SP1 was cloned in pCMV10-3XFLAG vector (Sigma). N-acetyl-leucinyl-leucinyl-norleucinal (ALLN) from Calbiochem. MG132 from Sigma.

Mutant phosphodegron was generated by site directed mutagenesis and primers were designed as mentioned in Quickchange primer designer tool. Sequences were verified by sequencing. Phosphodegron delta was cloned in pCMV10-3XFLAG vector. 3XFLAGβ-catenin, HA-S37A β-catenin, GST- N-terminus, GST-ARM repeats and GST-C-Terminus β-catenin were used as described (Notani et al., 2010). The siRNA sequences for shSP1 and shβ-catenin was designed using Dharmacon design centre. The siRNA sequence for siSP1, siβ-catenin and siGSK3β were procured from Dharmacon. All shRNAs were cloned in pSUPER Puro vector (OligoEngine). The details of siRNA and shRNA sequences are listed in Supplementary Table 1.

### RNA isolation and RT-PCR

RNA was isolated using Trizol reagent (Invitrogen). Two μg of RNA was used for first strand cDNA synthesis using Superscript III (Invitrogen). The cDNA was then used for quantitative PCR analysis in triplicates using an ABI 7500 Fast real-time PCR System (Applied Biosystems) as described (Ordinario et al., 2012). The sequences of oligonucleotide primers used for real-time quantitative PCR analysis are listed in Supplementary Table 2.

### Chromatin immunoprecipitation (ChIP) assay

ChIP assay was performed as described (Karmodiya et al., 2012). Briefly, cells were cross-linked by addition of formaldehyde to 1% final concentration in media and incubated at room temperature for 10 min, neutralized with 125 mM Glycine. Cells were then subjected to sonication using Covaris sonicator to fragment chromatin to obtain 200-500 bp fragments. Sonicated chromatin was precleared with nonsaturated beads. Precleared chromatin was incubated with specific antibodies and respective IgG types were used as isotype controls. Next day, beads saturated with tRNA and BSA were added (40 μl packed beads) and incubated for 4 h on rocker to pull down the antibody-bound chromatin and were subjected to elution using buffer containing SDS and sodium bicarbonate. Eluted chromatin was de-crosslinked and protein was removed by treating with proteinase K. Purified immunoprecipitated chromatin was subjected to PCR amplification using specific primers. Input chromatin was used as a control. ChIP primer details are listed in Supplementary Table 1.

### Immunoprecipitation, co-immunoprecipitation and ubiquitination

Immunoprecipitation and co-immunoprecipitation were done as essentially mentioned in (Cordenonsi et al., 2011). Briefly, cells were harvested and lysed in lysis buffer- 20 mM HEPES (pH 7.8), 400 mM KCl, 5% Glycerol, 5 mM EDTA, 0.4% NP40, phosphatase and protease inhibitors, lysates were sonicated and cleared by centrifugation. Before immunoprecipitation, lysates were diluted to 20 mM HEPES (pH 7.8), 50 mM KCl, 5% Glycerol, 2.5 mM MgCl_2_, 0.05% NP40 and incubated with the appropriate protein G-Dyna beads bound antibodies for four hours at 4°C (1/8^th^ to the lysis buffer). After three washes in binding buffer, co-purified proteins were analyzed by immunoblotting by using the ExactaCruz reagents (Santa Cruz Biotechnology) as secondary antibodies to reduce the background from IgG. Beads used for IP were pre-coated with antibody and then washed twice with binding buffer. In vivo ubiquitination was done as mentioned in Jeong et al 2012. Briefly cells were harvested and lysed in co-IP lysis buffer containing phosphatase, protease and de-ubiquitinase inhibitor NEM (N-ethylmaleimide, 10 mM) and IP and Co-IP were performed as mentioned above.

### GST pull-down

For GST pull-down, beads with purified proteins GST and GSTβ-catenin were incubated with lysate in binding buffer (25 mM HEPES (pH 7.9), 0.4 M KCl, 0.4% NP40, 5 mM EDTA, 1 mM DTT, 10% glycerol with protease inhibitors) for 1 h. After four washes with binding buffer, interaction of endogenous protein from cell lysate with purified proteins was determined by immunoblotting. For GST pull-downs with purified proteins FLAG-SP1 expressing cells, FLAG-SP1 was first immunoprecipitated using anti- FLAG antibody. After immunoprecipitation, FLAG-SP1 was eluted from beads using the FLAG peptide, followed by incubation with GST and GSTβ-catenin in binding buffer for 1h. After four washes with binding buffer, interaction complexes were resolved by SDS-PAGE and determined by immunoblotting. Domain Graph, version 1.0 was used to prepare protein domain structures (Ren J. et al, 2009).

### Cellular fractionation

For cytosolic and nuclear fractionation cells were washed in ice cold PBS. Cell pellets were then resuspended in hypotonic lysis buffer(10 mM Hepes, pH 7.9, 10 mM KCl, 0.1 mM EDTA, 0.1 mM EGTA 1 mM DTT) containing protease inhibitor and phosphatase inhibitor cocktail tablets. Cell suspensions were incubated on ice for 30 min to swell followed by adding 25 ul of 10% NP-40. Nuclear proteins, including the unlysed cells, were pelleted by spinning at 2,000 rpm for 2 min at 4_C centrifugation. The supernatant collected containing cytoplasm proteins. The pallet was then washed twice with hypotonic lysis buffer and then re-suspended in lysis buffer (20 mM Hepes, pH 7.9 0.4 M NaCl 1 mM EDTA 1 mM EGTA 1 mM DTT) containing protease inhibitor and phosphatase inhibitor cocktail tablets was then centrifuged at 14,000 rpm for 30 min at 4_C for nuclear fraction and analyzed further by SDS-PAGE followed by western blotting.

## Acknowledgements

Authors wish to thank Dr. Randall T Moon for gift of the TOP/FOP constructs. Work was supported by grants from the Centre of Excellence in Epigenetics program of the Department of Biotechnology, the Swarnajayanti Fellowship from the Department of Science and Technology, Government of India, and institutional funding from IISER Pune to SG. RM and AS are supported by fellowships from the University Grants Commission, India. SJP is supported by fellowship from the Council of Scientific and Industrial Research, Government of India.

## Author Contributions

RM and SG conceived project and designed experiments. Experiments were performed by RM, AS and SJP. SJP performed ChIP-seq data analysis. Manuscript is written by RM and SG.

